# Lack of evidence for cargo release of CD63-EVs into recipient cells

**DOI:** 10.1101/2025.04.01.646561

**Authors:** S. Askarian-Amiri, W. Weissenhorn, R. Sadoul, C. Chatellard

## Abstract

Extracellular vesicles (EVs), including exosomes, are widely believed to mediate intercellular communication by delivering molecular cargo to recipient cells. However, the efficiency and mechanisms of such cargo delivery remain unclear. In this study, we investigated whether EVs bearing the canonical exosomal marker CD63 are capable of fusing with recipient cell membranes to release their contents. Using the highly sensitive NanoBiT split luciferase system, we tracked potential fusion events between HiBiT-tagged CD63-EVs and HEK cells expressing cytosolic or endosome-localized LgBiT. While HiBiT and LgBiT reconstitution was readily detected when EVs carried the viral fusion protein VSV-G, no luminescence was observed with native CD63-EVs, despite their uptake or binding by recipient cells. Similarly, EVs carrying HiBiT-HSP70 failed to show evidence of cargo release. These findings demonstrate that unmodified CD63-EVs do not fuse with the plasma or endosomal membranes of recipient cells, suggesting that EV-mediated cargo delivery via membrane fusion is inefficient or absent under physiological conditions. Our results challenge the prevailing view of exosome function in intercellular communication and underscore the need for re-evaluation of EV cargo transfer mechanisms.

## Introduction

Cells secrete extracellular vesicles (EVs) either directly from the plasma membrane or through multivesicular endosomes that fuse with the plasma membrane ^1^. EVs allow secretion of biologically active molecules and are thought to allow cell-cell communication. Several studies have suggested that EV cargo can be delivered into recipient cells ^2 3^ though the efficiency of this process is generally considered as low ^4^. Furthermore, the molecular mechanisms underlying uptake and content delivery of EVs in acceptor cells remain poorly understood. To track exosome secretion and fate in recipient cells, many studies have utilized CD63 fused to GFP ^5 2 3^. Indeed, CD63 is a widely accepted exosome marker, highly enriched in the intraluminal vesicles of multivesicular endosomes which give rise to exosomes ^1,6^.

CD63-GFP-containing exosomes have been shown to bind to the surface of recipient cells, sometimes followed by endocytosis ^7 8^. In one study, CD63-labeled exosomes were reported to efficiently enter through endocytic hotspots and traffic within endosomes to sites of contact with the endoplasmic reticulum, where they were hypothesized to fuse and release their cargo ^9^. In this study, we aimed to demonstrate the potential fusion of CD63-bearing exosomes with HEK recipient cell membranes using the luminescence NanoBiT system.

The NanoBiT system is based on split luciferase complementation between two NanoLuc fragment: an 18-kDa polypeptide Large BiT (LgBiT), and an 11-amino-acid peptide, HiBiT. HiBiT binds to LgBiT with high affinity (Kd = 0.7 nM), thereby forming a stable complex that reconstitutes active NanoLuc luciferase. This system has previously been used to monitor the entry and release of HIV and virus-like particles containing HiBiT in cells expressing LgBiT ^10 11^. In our approach, we incubated EVs purified from cells expressing HiBiT fused to the N-terminal region of CD63, which is localized within the EV lumen (Fig.1c), with cells expressing cytoplasmic LgBiT. Fusion of exosomes with recipient cells should enable interaction between cytosolic LgBiT and HiBiT-CD63 thereby increasing luminescence. We found that such a luminescence increase did not occur with native exosomes and could only be detected with exosomes expressing the viral fusion protein VSV-G. The absence of detectable luminescence observed with unmodified exosomes indicates that fusion of CD63-containing-exosomes with receiving cells does not occur.

## Materials and Methods

### Cells Culture and Plasmids

The human embryonic kidney (HEK 293T) cell line was maintained in Dulbecco’s Modified Eagle Medium (DMEM) supplemented with 10% heat-inactivated fetal bovine serum (FBS) (Gibco). To generate a stable cell line expressing the LgBiT protein, HEK cells was transfected with the pCMV-LgBiT vector (Promega) using the calcium phosphate method. Twenty-four hours after transfection, the cells were treated with 200µg/µl hygromycin B to select for those expressing the LgBiT protein. HEK cells were incubated at 37°C in a humidified atmosphere with 5% CO_2_.

All cell lines were routinely tested for mycoplasma contamination using MycoAlert Mycoplasma Detection Kit (Lonza).

To generate the HiBiT-CD63-myc construct, HiBiT-Linker primers (Table 1) were paired and inserted into the hCD63-myc pcDNA3.1 vector using the BamH1 restriction site. Hsp70 cDNA was obtained from mouse brain cDNA via RT-PCR, using a forward primer incorporating the HiBiT sequence and a reverse primer containing the myc sequence (Table 1). The resulting HiBiT-Hsp70-myc cDNA was inserted into the Nhe1/Xba1 sites of the LgBiT expression vector (pCMV-LgBiT, Promega), to remove LgBiT cDNA and generating the pCMV-HiBiT-mHsp70-myc plasmid.

**Table 1:**
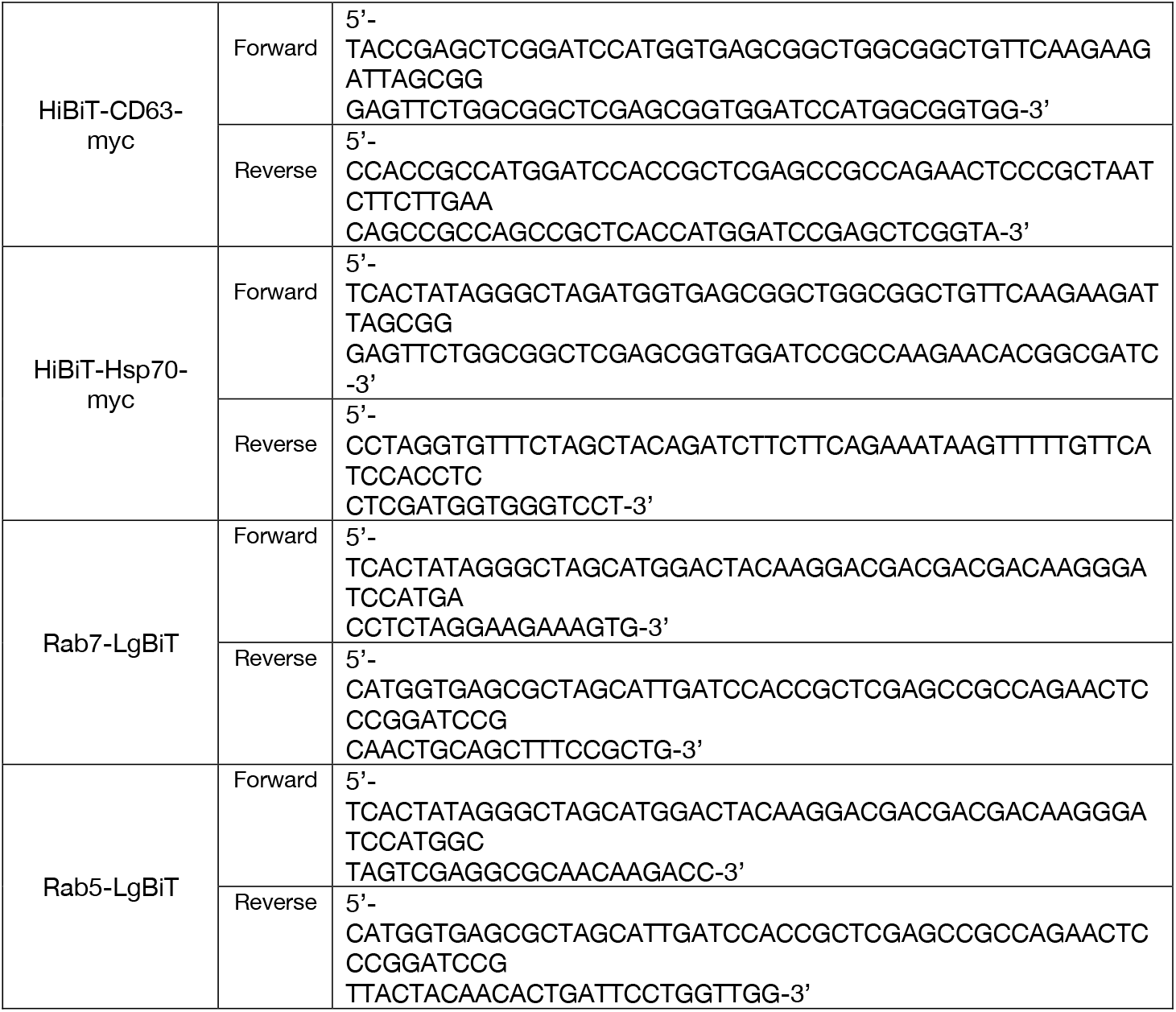
Primers used to generate plasmids.

hRab5-LgBiT and dRab7-LgBiT were constructed by amplifying (Table 1) and inserting hRab5 or dRab7 cDNA at the N-terminus of LgBiT into the Nhe1 site of the pCMV-LgBiT vector. The pCMV-VSV-G plasmid was purchased from Addgene (#8454). All plasmids were transformed into Top10 *E. coli* and subsequently purified using the Endotoxin-Free Plasmid Midiprep Kit (Macherey-Nalgene).

### Transfection

Transient transfection of HEK cells was performed using the calcium phosphate method. Briefly, 24 h prior to transfection, cells were seeded at a density of 4 × 10^4^ cells/cm^2^ on poly-ornithine-coated plates in culture medium. For co-transfections, the DNA ratio was 1:2 for HiBiT-CD63 to HiBiT-Hsp70, and 1:4 for VSV-G to HiBiT-CD63. Plasmids encoding the proteins of interest were diluted in sterile water and a CaCl_2_ solution (0.26 M final concentration), then mixed with an equal volume of 2× HEPES Buffered Saline (Sigma-Aldrich), gently bubbled to mix, and incubated for 20 min. The resulting DNA solution was added dropwise to the plated cells. Six hours later, the medium was aspirated and replaced with fresh culture medium.

### EVs isolation

Extracellular vesicles (EVs) from HEK cells were purified by differential centrifugation of cell culture supernatants. Briefly, HEK cells were transfected, and 6 h later, the cells were washed twice with PBS and incubated for 40 h in exosome-free medium (DMEM with 10% FBS, ultracentrifuged at 140 000 g for 18 h). The conditioned medium from the HEK cells was then subjected to serial centrifugation to remove cells, debris, and large vesicles: first at 2 000 g for 10 min, followed by 20 000 g for 30 min, collecting the supernatant and discarding the pellets. The medium was subsequently passed through a 0.22 µm sterile filter (Millex GV PVDF, Millipore). EVs were collected by ultracentrifugation at 100 000 g for 1.5 h and washed in filtered PBS before undergoing a second centrifugation at 100 000 g for 1.5 h. The final EVs were resuspended in PBS or Opti-MEM and immediately used for NTA or applied to the receiving cells.

### Western Blot analysis of cell lysates and EVs

Cell lysates were collected simultaneously with the EV supernatants. Briefly, cell lysis was performed using a buffer containing 20 mM Tris-HCl (pH 7.5), 150 mM NaCl, 1 mM MgCl_2_, 1% Triton X-100, and a complete, EDTA-free protease inhibitor cocktail (Roche). The lysates were clarified by centrifugation at 16 000 g for 5 min at 4°C, and the supernatant was transferred to fresh tubes and frozen at −20°C. Protein concentration was determined using the bicinchoninic acid (BCA) assay (Pierce). For each sample, 10 µg of cell lysate or 1µg of particles (equivalent to 4 × 10? particles) were separated on SDS-polyacrylamide gels, and the proteins were transferred to PVDF membranes (Millipore). The membranes were blocked in TBS-T milk solution (TBS, 0.05% Tween 20, 5% non-fat milk). Primary antibodies used included anti-myc rabbit polyclonal antibody (1:2000, Proteintech), anti-Alix rabbit polyclonal antibody (1:1000, Covalab), and anti-VSV-G rabbit polyclonal antibody (1:2000, Sigma-Aldrich). The membranes were then incubated with anti-rabbit IgG conjugated to HRP (1:10 000, Jackson ImmunoResearch) in TBS-T-milk solution for 1 h at room temperature. Immunoreactive bands were visualized using a solution containing 100 mM Tris-HCl (pH 8.5), 1.25 mM luminol, 2 mM p-coumaric acid, and 0.009% H_2_O_2_, and detected using the ChemiDoc MP Imager system (Biorad).

### Post Nuclear Supernatant of cells expressing LgBiT

Cells expressing LgBiT were washed two times with PBS, incubated within 20 mM Tris-HCl pH 7.5, 150 mM NaCl, 1 mM MgCl^2^ and gently detached from the plate using a scrapper. Cells were disrupted by sonication and lysate was centrifuged for 10 min at 1500 g. The supernatant was then ultracentrifuged at 100 000 g for 1 h and the supernatant was recovered as the cytosolic fraction.

### Nanoparticle Tracking Analysis (NTA)

The size and concentration distribution of EVs were measured using a NanoSight NS300 equipped with a 488 nm laser (Malvern Panalytical, Malvern, United Kingdom). EV samples were diluted 10- to 1 000-fold in filtered PBS to achieve a recording of 10–100 particles per frame. The movement of EVs was captured in three separate 60-second videos at camera level 15, and analyzed using NTA3.2 software (Malvern Panalytical). The data presented were the averages obtained from the analysis of three 60 s videos for each sample.

### EVs uptake

Receiving HEK cells were transfected with plasmids encoding LgBiT proteins one day after plating. Six hours post-transfection, cells were trypsinized and seeded at a density of 1 × 10^4^ cells/cm^2^ into poly-ornithine-coated white flat-bottom 96-well plates (ThermoScientific), then incubated for 24 h. Fresh EVs purified from cells expressing a HiBiT-fused protein were added to the receiving cells at 2 × 10^9^ particles per well and incubated at 37°C for the specified time period. At the indicated time, cells were washed twice with PBS and incubated with Opti-MEM, and when indicated, with 0.1 µM DrkBiT for 10 min at 37°C. Nano-Glo substrate (Nano-Glo Live Cell Assay System; Promega) was then added according to the manufacturer’s instructions and incubated for 20 min at 37°C. In some experiments, EVs were incubated in the presence of Nano-Glo substrate. Luminescence activity was measured using a ClarioStar microplate reader (BMG LabTech). After luminescence reading, 0.01% digitonin or 0.5% TritonX-100 were added to the same well, incubated 15 min at 37°C. Luminescence activity was measured under the same condition as above.

### Single vesicle imaging

EVs from HiBiT-CD63 /VSV-G double-transfected HEK cells were spotted onto coverslips previously ionized by plasma technology and left bound for 20 min. The EVs were incubated 20 min in 4% PFA solution, washed 3 times with PBS and then saturated using 5% BSA and 5% goat pre-immune serum. Immunostaining was performed using anti-CD63 monoclonal antibody (clone H5C6, BD Pharmigen) and anti-VSV-G rabbit polyclonal antibody and then revealed with anti-mouse IgG Alexa488 and anti-rabbit IgG Alexa594 secondary antibodies (Invitrogen). The coverslips were washed and mounted in Mowiol (Calbiochem) for observation using a 100 X oil objective of an IX83 inverted microscope (Olympus).

### Data analysis/statistical analysis

Statistical analysis was performed using GraphPad Prism 10 (GraphPad Software, San Diego, CA) with statistical significance considered at *p* ≤ 0.05. Statistical significance was assessed for non-parametric data by Kruskal-Wallis’s test with a post-hoc Dunn’s multiple comparisons test (unpaired, two-tailed) (Fig 3a, 4d), Mann-Whitney’s U test (unpaired, two-tailed) (Fig 2a) or Wilcoxon’s test (paired, two-tailed) (Fig 3a). For parametric data, normality of data was assessed using Shapiro-Wilk test then statistical significance was assessed by Student’s t test (paired, one-tailed) (Fig 1c, 2b).

**Fig. 1:**
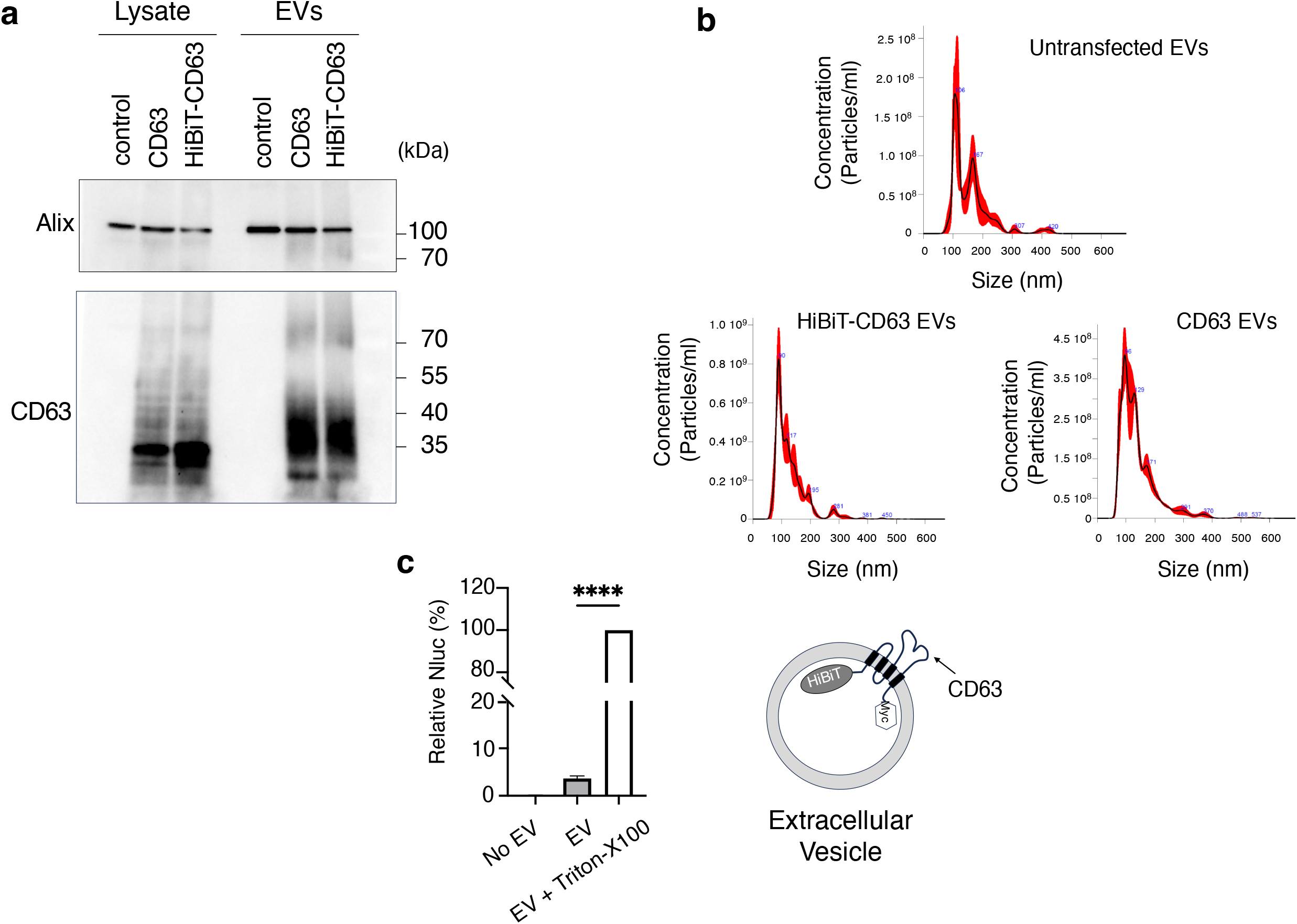
HiBiT-CD63 EV characterization. (a) CD63 and HiBiT-CD63 are enriched in EVs secreted by transfected HEK cells. 5µg of protein cell lysate and 1µg of EV proteins (4×10^9^ particles) were run on SDS-PAGE and blotted. CD63/ HiBiT-CD63 were revealed using anti-myc polyclonal antibody, while Alix was revealed using an anti-Alix polyclonal antibody. (b) The size and concentration distribution of EVs isolated from transfected cells do not differ from non-transfected cells as determined by NTA. The black line represents mean values of 3 videos, and the red shade the standard deviation. (c) The 11-amino-acid peptide, HiBiT, and a Myc tag were fused to the N-terminal and C-terminal region of CD63, respectively. Both tags are located within the EV lumen. EVs were incubated with the cytosol of LgBiT expressing cells, with or without 0.5% of Triton-X100. Luminescence of detergent solubilized EVs was set to 100%. Mean +/-SD of five independent wells from one representative experiment. Statistically significant difference was calculated using Student’s t test paired, two-tailed (****, p<0.0001).

**Fig. 2:**
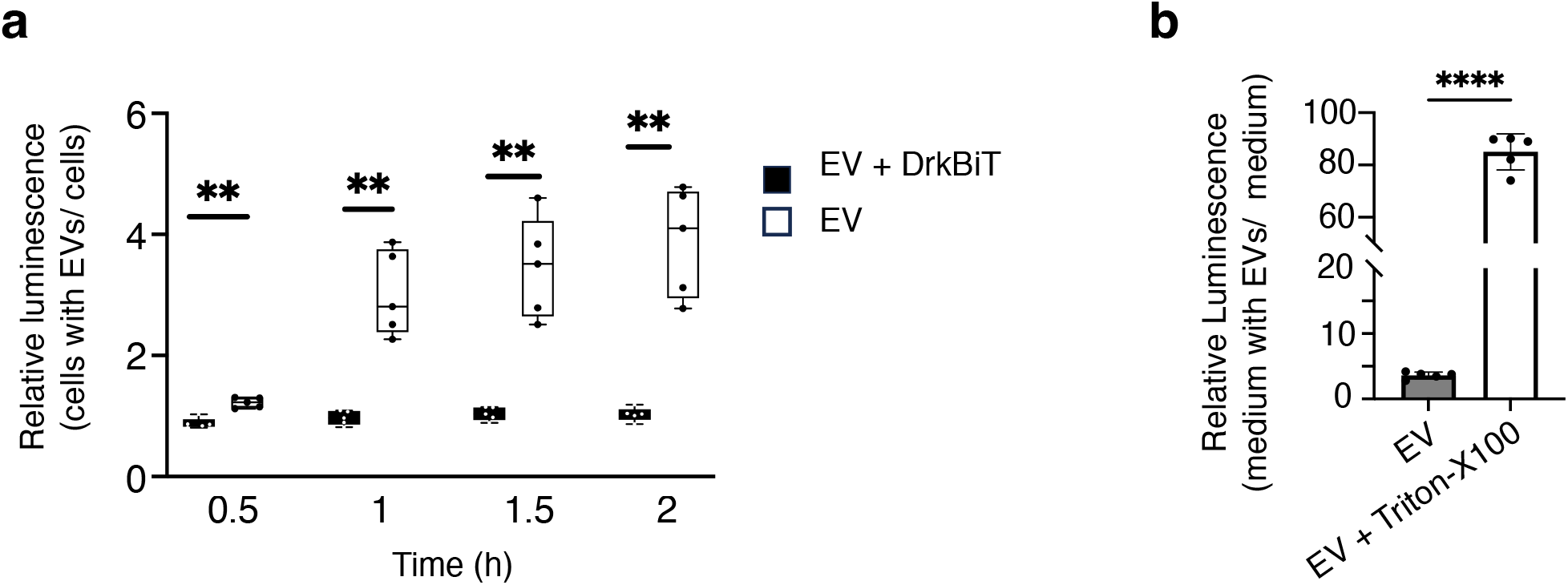
Luminescence increase over time is caused by LgBiT from the culture medium of LgBiT cells and is blocked by DktBiT. (a) HiBiT-CD63 EVs were incubated for the indicated times on LgBiT-expressing-HEK cells. At all times tested (0, 0.5, 1, 1.5 h), Nano-Glo substrate and 0.1µM of DrkBiT peptide were added together with EVs and luminescence measured 30 min later. Median (q1; q3) of five wells from one representative experiment. Statistically significant differences were calculated using Mann-Whitney’s U test (** *p* < 0.01). (b) Luminescence was measured in cleared medium of LgBiT expressing cells alone (no EV) or added of HiBiT-CD63 EVs incubated in absence or presence of 0.5% of Triton-X100. Mean +/-SD of five independent wells from one representative experimen. Statistically significant differences were calculated using Student’s t test paired, two-tailed (****, p<0.0001).

**Fig. 3:**
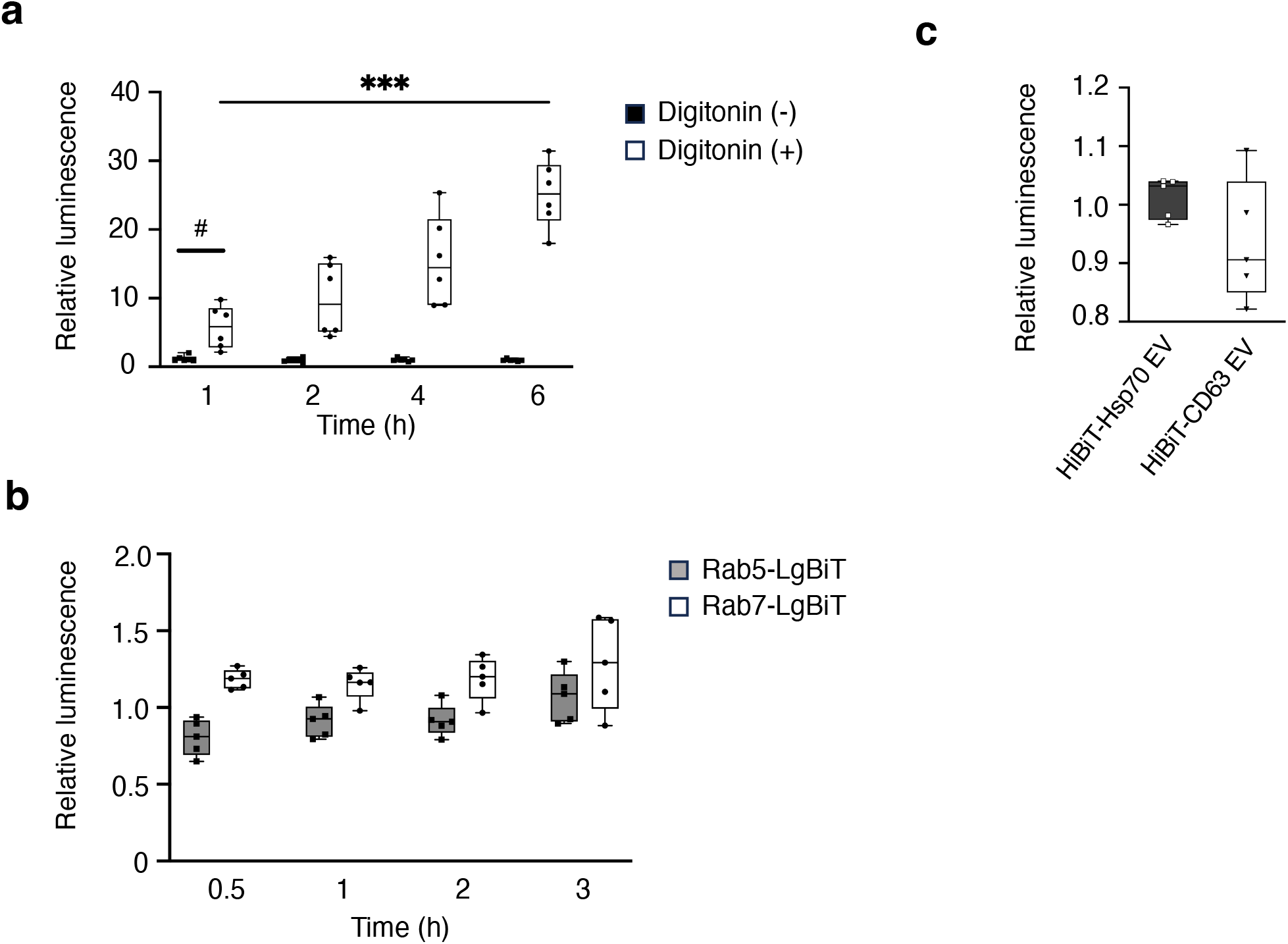
Binding and uptake of HiBiT-CD63 EVs by HEK cells. (a) HiBiT-CD63 EVs were incubated with LgBiT expressing HEK cells. At the indicated times, cells were washed and luminescence activity measured 30 min after addition of DrkBiT and Nano-Glo substrate. Relative luminescence is shown before (black box) and after (white box) addition of digitonin to induce LgBiT leakage from cells and access to HiBiT contained in the lumen of cell-bound EVs. Median (q1;q3) of three wells from two independent experiments. Statistically significant differences were calculated using Kruskal-Walli’s test (*** *p* < 0.001) and Wilcoxon’s test (# p<0.05). (b) : Incubation of EVs on cells expressing Rab5-, or Rab7-LgBiT does not indicate fusion of HiBiT-CD63 EVs. HiBiT-CD63 EVs were incubated on Rab5-LgBiT HEK cells (grey box) or Rab7-LgBiT HEK cells (white box). Cells were preincubated with DrkBiT for 10 min before addition of the Nano-Glo substrate together with DrkBiT and EVs. Luminescence was quantified after the indicated times in five and four independent wells for Rab5 and Rab7, respectively. Median (q1; q3) from one representative experiment is shown. (c) HiBiT-Hsp70 containing EVs do not fuse with LgBiT expressing cells. Luminescence activity of LgBiT-expressing cells was measured after a 3 h incubation of EVs containing HiBiT-Hsp70 (black box) or HiBiT-CD63 (white box) with LgBiT-expressing cells. The DrkBiT peptide was added 10 min before the Nano-Glo substrate. Median (q1; q3) of five wells from one representative experiment.

## Results

### Characterization of EVs containing HiBit-CD63

Extracellular vesicles (EVs) secreted by HEK cells expressing either HiBiT-CD63-myc or CD63-myc were isolated using ultracentrifugation. Western blot analysis revealed a significant enrichment of both HiBiT-CD63 and CD63 in the isolated EVs (Fig. 1a). NTA analysis demonstrated no notable alteration in the size of EVs secreted by HiBiT-CD63 cells compared to control (Fig. 1b). However, in line with previous findings by Sung et al. (2020) ^12^, CD63 overexpression appeared to increase the number of the smallest secreted EVs.

To verify the correct orientation of CD63 and the integrity of EVs purified by ultracentrifugation, we incubated HiBiT-CD63 EVs with LgBiT contained in cytosolic fractions prepared from LgBiT-expressing cells. After substrate addition, the recorded luminescence was less than 2% of that measured in the presence of Triton-X100, indicating that over 98% of HiBiT is enclosed within the EVs (Fig. 1c). This percentage, which is likely even higher as Triton-X100 reduces NanoLuc-induced luminescence ^13^ shows that the vast majority of HiBit fused to the N-terminal part of CD63 points inside EVs suggesting a correct orientation of the protein in the vesicles. Therefore, HiBiT-CD63 EVs were considered similar to wild-type EVs and were used in subsequent experiments.

### Incubation of EVs containing HiBiT-CD63 on LgBiT-expressing HEK cells does not increase luminescence

We then incubated HiBiT-CD63 EVs with LgBiT-expressing HEK-cells. In the initial set of experiments, cells were exposed to EVs for increasing durations, followed by washing and a 30-minute incubation with the cell-permeant luciferase substrate, Nano-Glo. Luminescence increased progressively with incubation time (Fig. 2a). However, no increase was detected when the Nano-Glo substrate was applied in the presence of the DrkBiT peptide, a HiBiT variant with a single Arg-to-Ala substitution that binds to LgBiT but does not restore NanoLuc activity (Fig. 2a). DrkBiT being membrane impermeable ^11^, the luminescence detected in absence of DrkBit is likely to reflect cell bound EVs with HiBiT exposed on the outside interacting with LgBiT released by receiving cells during incubation with the Nano-Glo substrate. Accordingly, luminescence was detected upon incubation of HiBiT-CD63 EVs with centrifuged culture media of LgBiT-expressing cells (Fig.2b), suggesting leakage of LgBiT from receiving cells. As expected, Triton-X100 lysis of the EVs, which exposed the luminal HiBiT, resulted in a several-fold increase in luminescence. The increase in luminescence correlating with the incubation time of vesicles on cells therefore likely reflects the increase in the number of vesicles bound to the surface of recipient cells.

Next, LgBiT-HEK cells were incubated with HiBiT-CD63 EVs for increasing time periods, alongside a low amount of DrkBiT to block any luminescence caused by LgBiT leakage during the incubation. EVs that bound to, or were endocytosed by LgBiT-expressing HEK cells were identified by the luminescence detected after addition of a low concentration of digitonin to permeabilize both cells and EVs. As shown in Fig. 3a, there was a significant time-dependent increase in luminescence only seen with digitonin. This suggests that EVs bind to and are possibly endocytosed by LgBiT-expressing cells but do not undergo the membrane fusion step required for the interaction between LgBiT and HiBiT.

All of the above experiments were conducted with LgBiT expressed in the cytosol of recipient cells. Since EVs have been suggested to fuse with endosomes ^9^, we next explored the use of LgBiT fused to proteins located on the cytosolic surface of endosomes. HiBiT-CD63 EVs were incubated with HEK cells expressing LgBiT fused to either Rab5 or Rab7 (Fig. 3b), proteins associated with early and late endosomes, respectively. However, even in this case no fusion of CD63-bearing EVs with endosomal membranes could be demonstrated since luminescence remained near background levels at all times tested.

Finally, we also used HiBiT fused to HSP70, which has been shown to concentrate inside EVs ^14 3^ to monitor cytosolic release of EV cargoes.

Fig. 3c shows that, as in the case of HiBiT-CD63, no sign of increase in luminescence could be detected upon incubation of HiBiT-HSP70 containing EVs with LgBiT expressing cells.

Thus, our results suggest that despite EVs binding and possibly endocytosis by recipient cells, EVs do not fuse with membranes of the receiving cells, as evidenced by the lack of HiBiT-CD63 or HiBiT-HSP70 interaction with cytosolic or endosome bound LgBiT.

### CD63-EV cargo release into receiving cells can be measured with EVs carrying VSV-G

To confirm that the split NanoLuc assay was sufficiently sensitive to detect fusion events of EVs with membranes of receiving cells, we purified EVs from HEK cells co-expressing the VSV-G fusion protein with HiBiT-CD63. NTA analysis revealed that the total number of EVs was comparable between HEK cells expressing HiBiT-CD63 alone and those co-expressing HiBiT-CD63 with VSVG (Fig. 4a left panel), even if the size of EVs from VSV-G expressing cells was slightly increased (Fig.4a right panel).

**Fig. 4:**
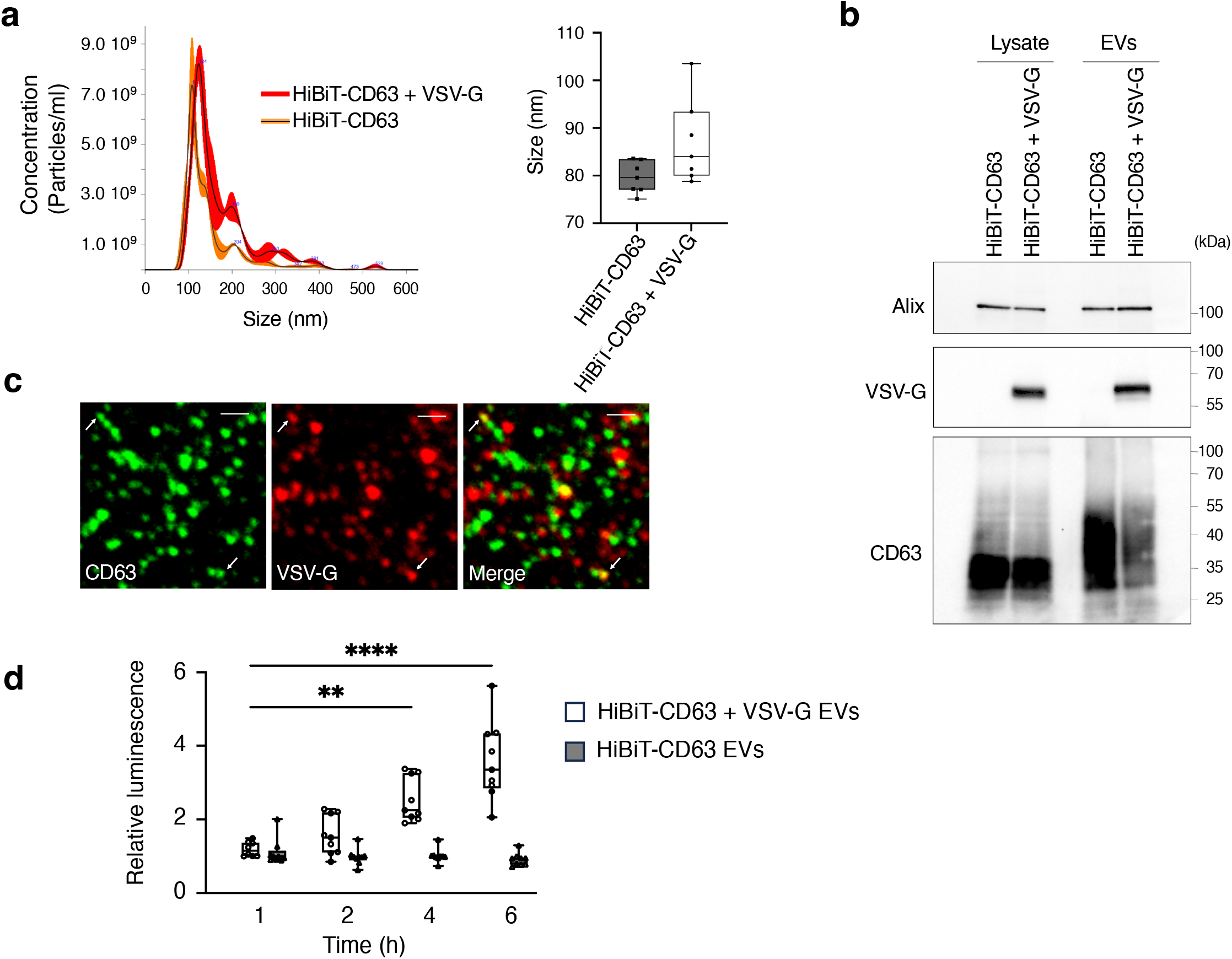
Fusion of HiBiT-CD63 EVs containing VSV-G protein can be detected. (a) Left panel: NTA analysis of HiBiT-CD63 EVs and HiBiT-CD63+VSV-G EVs. The black line represents mean values of three videos and the red shade represents the standard deviation. Right panel: The particle size from NTA was measured and compared between both groups. Median (q1; q3) of seven independent experiments. (b) VSV-G is incorporated in EVs purified from cells expressing both HiBiT-CD63 and VSV-G. Western blot analysis of lysates and EVs secreted by cells transfected with HiBiT-CD63 or HiBiT-CD63 together with VSV-G. (c) VSV-G protein can be detected in some EVs containing CD63. EVs purified from HiBiT-CD63/VSV-G double-transfected HEK cells were spotted onto coverslips, stained with anti-CD63 monoclonal (green) and anti-VSV-G polyclonal (red) antibodies, and then imaged using confocal fluorescence microscopy. Arrows indicate double-labelled EVs. (Scale bar = 1µm). (d) HiBiT-CD63 + VSV-G EVs fuse with membranes of LgBiT-HEK cells. EVs were incubated on LgBiT-cells and washed at the indicated times. DrkBiT was added 10min before the Nano-Glo substrate and luminescence measured 20 min later. Median (q1; q3) of three wells from three independent experiments. Statistically significant differences were calculated using Kruskal-Walli’s test (** p< 0.01, **** *p* < 0.0001).

WB analysis showed the presence of VSV-G in EVs purified from cells expressing both HiBiT-CD63 and VSV-G (Fig. 4b). We noticed that the amount of HiBiT-CD63 is significantly decreased in EVs secreted by cells also expressing VSV-G. Co-staining of EVs purified from cells expressing CD63 and VSV-G (Fig. 4c) showed that only a small fraction of the CD63 exosomes carried VSV-G as already shown by Zhang et al. (2022) ^*15*^. However, this minimal amount of VSV-G was sufficient to induce fusion of CD63 exosomes. Indeed, luminescence measured in LgBiT cells incubated with EVs purified from cells expressing both HiBiT-CD63 and VSV-G increased within 2 hours, whereas no change was seen with EVs from cells expressing only HiBiT-CD63 (Fig. 4d) even with incubation up to 18h (not shown). Thus, the NanoBiT system is sensitive enough to monitor fusion of HiBiT-CD63 EVs to membranes of LgBiT expressing cells if EVs carry the fusion protein VSV-G.

## Discussion

Our study set out to employ the NanoBiT assay to assess whether extracellular vesicles (EVs) bearing HiBiT-CD63 could fuse with HEK recipient cells expressing LgBiT. The NanoBiT system, which relies on the high-affinity interaction between HiBiT and LgBiT (Kd ≈ 0.7 nM), was chosen for its remarkable sensitivity and specificity. This design ideally enables the detection of even rare fusion events by reconstituting an active luciferase enzyme upon membrane fusion. Our results demonstrate that the assay is indeed highly sensitive. When EVs containing HiBiT-CD63 were incubated with LgBiT-expressing cells, a robust and rapid increase in luminescence was evident when EVs were engineered to express the viral fusion protein VSV-G. This clear positive control confirms that the NanoBiT system is highly sensitive in reporting fusion events when they occur. The high potency of our system to report fusion is reinforced by the fact that VSV-G was present only in a low number of CD63 exosomes consistent with the observation of Zhang et al reporting that less than 5% of CD63-EVs incorporate VSV-G-GFP ^15^. EVs containing HiBiT-CD63 but lacking VSV-G bound to and were possibly endocytosed by receiving cells but led to no increase in luminescence showing that in the absence of any fusogenic stimulus, the assay detects little to no membrane fusion between EVs and cell membranes.

Moreover, the use of the DrkBiT peptide to block signals arising from LgBiT leakage further attests to the assay’s specificity, ensuring that any luminescent readout is attributable solely to the fusion-mediated mixing of the HiBiT and LgBiT components. We also fused LgBiT to endosomal markers (Rab5 and Rab7) in order to concentrate it to endosomal membranes and thereby increasing the chance of interaction of HiBiT-CD63 with LgBiT. Here again, no increase of luminescence suggesting fusion of exosomes to the limiting membrane of endosomes could be observed. Similarly, changing CD63 for HSP70 as a reporter of EV cargo release failed to generate a luminescence signal when incubated on LgBiT expressing cells. Similarly, Somiya et al failed to detect luminescence in LgBiT expressing cells incubated with EVs containing HiBiT fused to a de novo designed protein (I3-01) that spontaneously forms a 60-mer self-assembled nanocage (EPN-01) ^16^. Here again, a time increase in luminescence could only be detected when EPN-01 containing EVs also contained the fusion protein VSV-G ^17^. Collectively, our findings imply that natural, unmodified CD63-exosomes bind to cells and might primarily be internalized via endocytic pathways without significant membrane fusion—a scenario that could limit the functional delivery of their cargo under physiological conditions. Noteworthy, is that EPN-01 containing EVs ^17^ were shown to be secreted from the plasma membrane ^16^ implying that the lack of fusion of EVs with recipient cells is not specific for exosomes but concerns also ectosomes.

Since we and others ^8,9^ have demonstrated that endocytosis of CD63-exosomes occurs, it seems surprising that these exosomes are not capable of fusing with the limiting membrane of endosomes unless they contain VSV-G. Indeed, a chemically tunable cell-based system allowed Perrin et al. (2021) to demonstrate that in endosomes, commitment to intraluminal vesicles (ILVs) carrying CD-63 is not a terminal event, and that a return pathway exists, allowing ‘‘back-fusion’’ of intraluminal membranes to the limiting membrane of multivesicular bodies ^18^. However, they also demonstrate that only a pool of ILVs contributes to dynamics within MVBs allowing intraluminal proteins to return to the limiting membrane, whereas the other, more inert pool encompasses the bulk of secreted exosomes. This might explain why exosomes are not capable of fusing with endosome limiting membranes of receiving cells.

In conclusion, our study demonstrates that the NanoBiT assay is a sensitive and specific method for detecting EV membrane fusion. The lack of detectable fusion from unmodified exosomes, contrasted with the strong signal observed from VSV-G-modified EVs, suggests that native exosomes may not efficiently fuse with recipient cell membranes under the tested conditions. These findings highlight the need for further investigation into the fusion capabilities of natural EVs. Alongside growing evidence pointing to the limited ability of EVs to deliver functional cargo to recipient cells ^2-4,17,19^, our results call into question the widely held view that EVs are key mediators of intercellular communication through cargo transfer.

## Acknowledgments

We thank B. Blot for helping with some of the cDNA constructs, M.O. Fauvarque, J. Fauré, S. Fraboulet for helpful discussions throughout this work. We also thank K. Sadoul for critically reading the manuscript.

This work was financed by ANR labex (Grenoble Alliance for Integrated Structural Cell Biology). The funders had no role in study design, data collection and analysis, decision to publish, or preparation of the manuscript Competing interests: The authors have declared that no competing interests exist.

## Author contributions

A.A.S. performed the EV purification and characterization (NTA), did most of NanoLuc assays, made DNA constructions, W.W. and R.S. funding acquisition, R.S. supervised the project and wrote the manuscript, C.C. performed all experiments with VSV-G-EVs, made statistical analysis and co-supervised the project. A.A.S. and C.C. writing original draft preparation.

